# Sarcopenia is an independent predictor of hospitalization in chronic kidney disease outpatients

**DOI:** 10.1101/516096

**Authors:** Hye Yun Jeong, Wooyeol Ahn, Jun Chul Kim, Yu Bum Choi, Jinkwon Kim, Hak Hoon Jun, Soonchul Lee, Dong Ho Yang, Jisu Oh, Jinkun Bae, So-Young Lee

## Abstract

**Background:** Patients with chronic kidney disease (CKD) experience much more marked and earlier muscle wasting than subjects who do not have chronic illnesses. However, a few studies that have examined sarcopenia have been reported in CKD patients. We investigated the prevalence of sarcopenia in predialysis and dialysis outpatients with CKD and explored its relationship with the clinical outcomes.

**Measurements:** Sarcopenia was defined as reduced muscle strength accompanied by decreased adjusted appendicular skeletal muscle (ASM), while those patients who exhibited only one of these characteristics were categorized as presarcopenic patients. ASM was measured by bioimpedence analysis, and muscle strength was evaluated by handgrips. ASM was adjusted by weight (ASM/wt). Patients were prospectively followed for up to 2 years.

**Results:** One hundred seventy-nine patients were recruited (114 male and 65 female patients who were classified into 103 predialysis patients and 76 dialysis patients, with 44.7% having diabetes). Their mean age was 60.6 ± 13.5 years old. The prevalence of sarcopenia was 9.5%, while 55.9% of the patients were categorized as presarcopenic. The ASM/wt index showed significant correlations with age, handgrip strength, HOMA-IR and frailty scores. Multivariate Cox proportional hazards models demonstrated that the risk of hospitalization was significantly higher for patients with presarcopenia [hazard ratio (HR), 2.48; 95% confidence interval (CI), 1.180–5.230], and the risk of hospitalization was much higher for patients with sarcopenia than for patients in the nonsarcopenic group (HR, 9.11; 95% CI, 2.295–25.182)

**Conclusions:** Sarcopenia and presarcopenia, which were defined using the ASM/wt index and handgrip strength, predicted a poorer, hospitalization-free survival in CKD patients

## Introduction

Sarcopenia is defined as the degenerative loss of skeletal muscle mass and strength associated with aging (1). However, patients with chronic disease experience much more marked and earlier muscle wasting than that of subjects who do not have chronic illnesses (2). Furthermore, sarcopenia not only has been associated with bone loss and a frail phenotype (3, 4) but also has been found to be a major contributor to poor clinical outcomes, including mortality, hospitalization, institutionalization, and disability (5-7).

Although often recognized as a comorbidity of hypertension or diabetes, chronic kidney disease (CKD) by itself contributes to global morbidity and mortality by increasing the risks related to cardiovascular diseases and infection (8, 9). In a report by the World Health Organization, CKD is represented as a single disease with a tremendous economic burden, of which more than 2-3% of the annual healthcare budget is required for the treatment of this disease in high-income countries (10). In this context, it is noteworthy that recent reports have observed close relations between either physical inactivity or low muscle mass and CKD (11-13). Moreover, exercise for muscle training has reportedly resulted in positive effects in this population of patients (14, 15), which suggests that the detection of muscle failure, along with the resulting appropriate intervention, could improve clinical prognoses in patients with CKD.

Although sarcopenia has been recognized as a disease (via the classification of muscle failure with an ICD-10 code that was established in 2016 (16)), the operational definitions for sarcopenia have long been undetermined, and studies examining the incidence of sarcopenia in patients with CKD are still limited.

In this study, we investigated the prevalence of sarcopenia in predialysis and dialysis outpatients with chronic kidney disease (CKD) according to the Asian Working Group for Sarcopenia (AWGS) recommendation and explored its relationship with clinical outcomes.

## Methods

### Patients

One hundred three patients with CKD and 76 patients with ESRD on maintenance hemodialysis were recruited and prospectively followed for up to 2 years. The criteria for inclusion in this study included patients who were older than 20 years old with a confirmed diagnosis of CKD [defined as patients who were on dialysis or who had 2 previously estimated glomerular filtration rate (eGFR) values < 60 mL/min/1.73 m^2^, which was calculated according to the equation of the Modification of Diet in Renal Disease Study Group and was obtained at an interval of 3-6 months]. Patients were categorized according to CKD stages and Kidney Disease Outcomes Quality Initiative guidelines (17). Patients who were categorized as CKD Stage 5D were on hemodialysis 3 times/wk (> 12 h/wk) for at least 3 months without renal transplantations. All of the participants provided prior written consents. No patients had a history of cancer, coagulation disorders, or active infection. Any patients who were unable to ambulate, either with or without assistive devices, or had insufficient cognitive function to communicate with the interviewer were excluded. The study was approved by the Institutional Review Board of our Medical Center.

### Grip Strength and Physical Performance

The patients performed three tests of maximum handgrip strength with a Jamar hand dynamometer (Sammons Preston Inc., Bolingbrook, IL). Low handgrip strength was defined as < 26 kg for men and < 18 kg for women, according to the AWGS recommendation (18). Slow walking speed was measured based on the time to walk 4 m, and the cutoff value for low gait speed was ≤ 0.8 m/s, as suggested by the AWGS (18).

### Skeletal muscle mass measurement

Height was measured by using a stadiometer. The postdialysis weights were recorded from the last three dialysis sessions, and the average of these weights was calculated in the patients undergoing hemodialysis. To assess body composition, we used a bioimpedance analysis machine (Inbody 620, In-body, Seoul, Korea) with measuring frequencies of 5, 50, and 500 kHz. Weight-adjusted, squared height-adjusted, and body mass index (BMI)-adjusted appendicular skeletal muscle (ASM) was assessed in all of the subjects. Decreased ASM was defined as a weight-adjusted ASM (ASM/kg*100) less than 32.2% for men and less than 25.6% for women (19), a squared height-adjusted ASM (ASM/ht^2^) less than 7.0 (kg/m^2^) for men and less than 5.7 (kg/m^2^) for women (20), or a BMI-adjusted ASM (ASM/BMI) less than 0.789 (m^2^) for men and less than 0.512 (m^2^) for women (21).

### Definition of sarcopenia

Sarcopenia was considered to be present when subjects had low handgrip strengths accompanied by a low adjusted ASM. Those subjects who showed low handgrip strengths or low muscle volumes were categorized as being presarcopenic (22).

### Definition of frailty

We adopted the Fried criteria as the definition of frailty (23). At enrollment, the five components of this frailty scale were measured: shrinking (a self-report of unintentional weight loss of more than 10 pounds in the past year based on dry weight, i.e., the weight of an individual undergoing hemodialysis without the excess fluid that builds up between dialysis treatments, which is more representative of the weight that subjects would have if they had normal kidney function), weakness (a grip-strength below an established cutoff value, based on sex), exhaustion (a self-report), low activity (kcal/wk values below an established cutoff value), and slow walking speed (the time taken to walk 4 m below an established cutoff value, according to sex) (23). A score of 1 was given for each of the measured components. The aggregate frailty score was calculated as the sum of the component scores (range 0–5), and subjects were categorized as frail if the patients received 3 or more points (24).

### Clinical variables

The patient demographic and clinical data, including age, sex, etiology of CKD (e.g., diabetes, hypertension, glomerulonephritis, polycystic kidney disease or unknown disease) and other comorbidities, were obtained via medical record reviews. Cardiac diseases were defined as patients with any medical histories of angina pectoris, a positive treadmill test, myocardial infarction, percutaneous transluminal coronary angioplasty, coronary artery bypass surgery, or congestive heart failure. Cerebrovascular diseases were defined as patients with medical histories of a stroke, a transient ischemic attack, or an intracranial hemorrhage. Laboratory findings were collected, including serum hemoglobin, serum calcium, blood urea nitrogen, phosphate, intact parathyroid hormone (iPTH), uric acid, total cholesterol, low-density lipid (LDL) cholesterol, c-reactive protein (CRP), 25-hydroxyvitamin D (25[OH]D), and albumin levels at the time of patient enrollment. The Homeostatic Model Assessment for Insulin Resistance (HOMA-IR) was calculated according to the following formula: [fasting insulin (μU/L)*fasting glucose (nmol/L)]/22.5 (25).

### Clinical outcome: hospitalization-free survival

We prospectively observed all hospitalization events, mortalities, and kidney transplantations over a 2-year follow-up period. A hospitalization was defined as any hospitalization, regardless of the reason for admission, with more than 1 overnight stay. The hospitalization causes were classified as cardiac and/or cerebrovascular, infectious, or other causes via medical record reviews or telephone contacts. The outcome for this analysis was time to hospitalization from any cause.

### Statistical analysis

The categorical variables were recorded as numbers and percentages. The continuous variables are presented as the mean ± standard variation or median (IQR). Student’s t-tests, Mann-Whitney U tests or ANOVAs were used to compare the continuous variables. The categorical variables were compared using χ2 tests or Fisher’s exact tests. Pearson’s correlation coefficients were used to summarize the cross-sectional relationships among age, hand grip strength, HOMA-IR, and ASM. Kaplan-Meier curves were used to estimate event times, and the distributions were compared via log-rank tests. A Cox regression model was used to analyze the independent variables that were associated with hospitalization or mortality. A p-value < 0.05 was considered to be statistically significant. Statistical analyses were performed using SPSS for Windows (version 21; SPSS, Chicago, IL, USA).

## Results

### Study population

Table 1 shows the baseline clinical and biochemical characteristics of the study population. The mean age was 60.6 ± 13.5 years old. Of the patients in this study, 63.6% were male, and 44.7% had diabetes. The mean estimated glomerular filtration rate was 39.2 ± 20.4 ml/min/1.73 m^2^ among the predialysis patients. Of the predialysis patients, 28.9% were classified as Stage 3a, 30.1% were classified as Stage 3b, 24.1% were classified as Stage 4, and 16.9% were classified as Stage 5. Chronic hemodialysis patients (Stage 5D) comprised 42.5% of the patients in this study, and their mean duration of dialysis was 52.9 ± 47.7 months. The mean Kt/V of the chronic hemodialysis patients was 1.7 ± 0.3.

**Table 1.** Baseline characteristics

| Characteristics | Men<br>(n =114) | Women<br>(n =65) | P-value |
| --- | --- | --- | --- |
| Age, years | 60.9 ± 13.4 | 60.1 ± 13.8 | 0.705 |
| Dialysis, n (%) | 45 (39.5) | 31 (47.7) | 0.346 |
| HTN, n (%) | 92 (81.4) | 50 (76.9) | 0.474 |
| Diabetes, n (%) | 51 (44.7) | 29 (44.6) | 0.987 |
| Cerebrovascular, n (%) | 19 (16.7) | 4 (6.2) | 0.044 |
| Cardiac, n (%) | 18 (15.9) | 7 (10.8) | 0.341 |
| ALM, kg | 26.0 ± 5.4 | 17.1 ± 2.9 | < 0.001 |
| BMI, kg/m<sup>2</sup> | 23.0 ± 4.7 | 23.3 ± 5.0 | 0.669 |
| Height, m | 1.69 ± 0.07 | 1.55 ± 0.06 | < 0.001 |
| Weight, kg | 68.6 ± 15.4 | 55.1 ± 9.3 | < 0.001 |
| ECW/TBW, | 0.38 ± 0.02 | 0.38 ± 0.02 | 0.913 |
| Hand grip strength, kg | 24.8 ± 9.3 | 15.4 ± 5.6 | 0.008 |
| Walking speed, m/s | 0.98 ± 0.40 | 0.97 ± 0.41 | 0.885 |
| MAP, mmHg | 93.6 ± 22.0 | 89.2 ± 24.1 | 0.214 |
| HOMA-IR | 2.4 (1.4 - 5.7) | 2.4 (1.2 - 5.4) | 0.943 |
| 25(OH)D | 21.1 ± 16.7 | 13.3 ± 8.4 | <0.001 |
| Uric acid, mg/dL | 7.4 ± 2.1 | 7.4 ± 1.9 | 0.996 |
| Hemoglobin, g/dL | 12.1 ± 3.9 | 11.0 ± 1.5 | 0.022 |
| Creatinine, mg/dL | 5.6 ± 4.4 | 5.7 ± 4.3 | 0.932 |
| WBC, *10<sup>3</sup>/μL | 6.3 ± 2.1 | 6.3 ± 1.6 | 0.817 |
| Protein, g/dL | 6.7 ± 0.6 | 6.6 ± 0.6 | 0.460 |
| Albumin, mg/dL | 4.1 ± 0.5 | 4.7 ± 0.4 | 0.902 |
| Calcium, mg/dL | 8.8 ± 0.7 | 8.9 ± 0.6 | 0.656 |
| Phosphate, mg/dL | 4.3 ± 1.3 | 4.8 ± 1.4 | 0.002 |
| iPTH, mg/dL | 81.9 (40.5 - 144.7) | 89.3 (45.6 - 229.4) | 0.087 |
| T. cholesterol, µg/dL | 153.8 ± 47.8 | 161.8 ± 42.8 | 0.266 |
| LDL cholesterol, mg/dL | 84.4 ± 36.6 | 79.3 ± 33.0 | 0.379 |
| CRP, mg/dL | 0.07 (0.03 - 0.20) | 0.05 (0.03 - 0.13) | 0.872 |
| Glucose, mg/dL | 132.9 ± 57.9 | 141.2 ± 76.4 | 0.417 |

Data are presented as number (%), mean ± standard deviation, median (interquartile range).
25(OH)D, 25-hydroxyvitamin D; BMI, body mass index; CRP, c-reactive protein; ECW/TBW, extracellular water/total body water; HOMA-IR, Homeostatic Model Assessment for Insulin Resistance; HTN, hypertension; iPTH, intact parathyroid hormone; LDL cholesterol, low density lipid cholesterol; MAP, mean arterial blood pressure; T. cholesterol, total cholesterol; WBC, white blood cell.

### Prevalence of sarcopenia and associated factors

Based on the 3 different indices (ASM/wt, ASM/ht^2^ and ASM/BMI; Table 2), seventeen (9.5%), eight (4.5%), and five (2.8%) patients had sarcopenia, respectively, while 100 (55.9%), 103 (57.5%), and 105 (58.7%) patients were categorized as presarcopenic, respectively. Approximately 60% of the CKD patients had either sarcopenia or presarcopenia, according to all three of the ASM indices.

**Table 2.** Prevalence rate of sarcopenia in the patients with chronic kidney disease according to three different operational methods.

| **Sarcopenia characteristics** | **Weight based** | **Height based** | **BMI based** |
| --- | --- | --- | --- |
| Sarcopenia, n (%) | 17 (9.5) | 8 (4.5) | 5 (2.8) |
| Presarcopenia, n (%) | 100 (55.9) | 103 (57.5) | 105 (58.7) |
| Normal, n (%) | 62 (34.6) | 68 (38.0) | 69 (42.1) |

The analysis via the use of Pearson’s correlation coefficients revealed that the absolute value of ASM was negatively correlated with age (r = −0.432, p < 0.001) and frailty scores (r = −0.330, p < 0.001) but positively correlated with grip strength (r = 0.607, p < 0.001, Figure 1). The ASM/wt index had inverse correlations with age (r = - 0.247, p = 0.009), HOMA-IR (r = −0.202, p = 0.015) and frailty scores (r = −0.291, p < 0.001) and had a positive correlation with handgrip strength (r = 0.371, p = 0.009) (Figure 1). The ASM/Ht^2^ index was only modestly correlated with handgrip strength (r = 0.239, p = 0.002) and frailty scores (r = −0.261, p = 0.004). However, the ASM/BMI index showed no significant relationships with age, handgrip strength, HOMA-IR or frailty scores (Figure 1).

**Figure 1.**
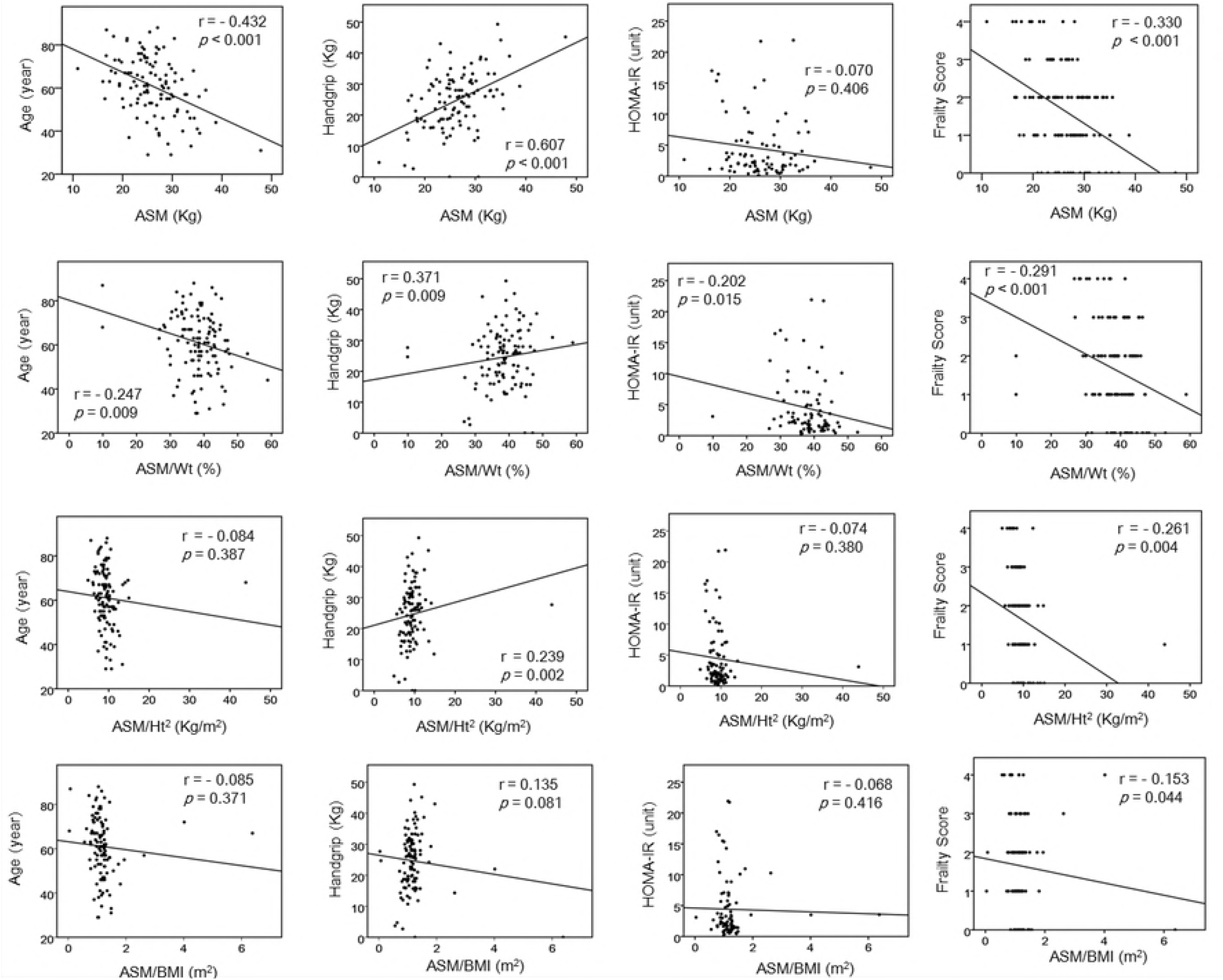
Scatterplots and correlations of appendicular skeletal muscle (ASM) or adjusted ASM indices versus age, handgrip strength, HOMA-IR or frailty scores for the patients with chronic kidney disease. ASM/BMI, body mass index adjusted ASM; ASM/ht2, height square adjusted ASM; ASM/wt, weight adjusted ASM; HOMA-IR, Homeostatic Model Assessment for Insulin Resistance; r, Pearson’s correlation coefficients; *p*, P-value.

### Characteristics based on sarcopenia status categorized by the ASM/wt index

The ASM/wt index showed the best correlations with chronological age, muscle strength, insulin resistance and geriatric syndrome, which suggests that the ASM/wt index might be the most appropriate and practical index in our subjects. In view of this contention, we divided our subjects into normal, presarcopenia, or sarcopenia groups, based on the ASM/wt index (Table 3).

**Table 3.**
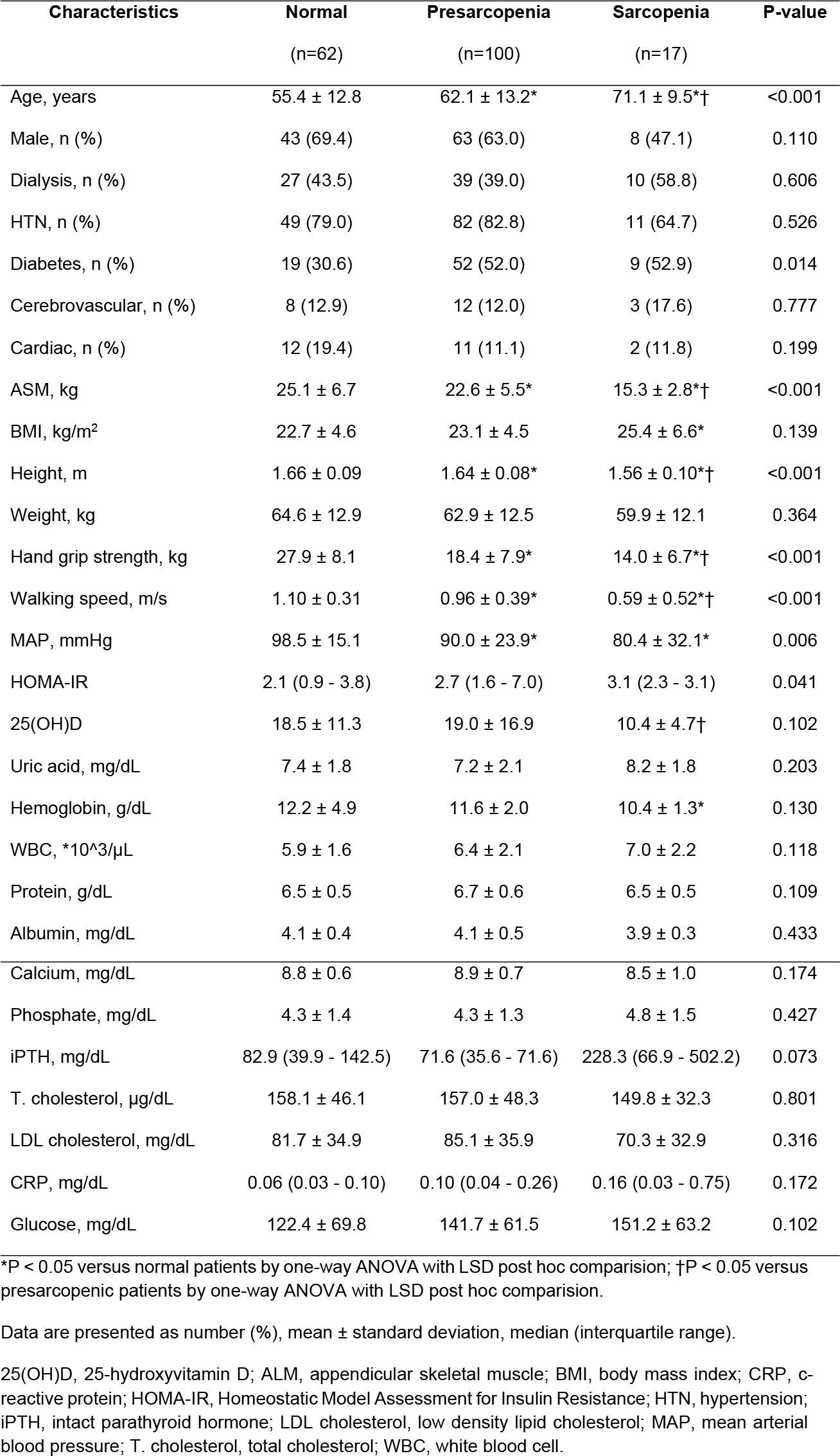
Comparison of clinical and laboratory characteristics according to sarcopenia status categorized by ASM/wt index.

The patients with presarcopenia were older than the normal patients, and the patients with sarcopenia were much older than the patients with presarcopenia (p < 0.001). More diabetic patients were included in the groups of presarcopenia and sarcopenia patients than were included in the normal patient group (p = 0.014). Handgrip strength and walking speed gradually decreased in the patients with presarcopenia and sarcopenia compared with those in the patients with normal skeletal muscle mass and function (p < 0.001). HOMA-IR was significantly and progressively increased in the presarcopenia and sarcopenia patients (p = 0.041), which indicates that they may have metabolic problems compared to those patients in the normal group. The patients with sarcopenia had relatively lower hemoglobin, albumin, calcium and 25(OH)D levels, but they were not statistically significantly lower. Sarcopenic patients had trends for higher phosphate, iPTH, and CRP levels, but these trends were not statistically significant.

### Sarcopenia and Clinical Outcomes

Fifty-one patients (25 predialysis and 26 dialysis patients) were hospitalized, 6 patients died (1 predialysis and 5 dialysis patients), and 2 patients underwent kidney transplantations during the 2-year observational period. The mean observational period was 552.0 ± 252.8 days. The causes of hospitalization events included cardiac and/or cerebrovascular disease (33.3%), infection-related disease (25.5%), and initiations of dialysis (23.5%). The most common causes of death were pneumonia (50%) and cardiac arrest (20%).

The hospitalization-free survival, according to the sarcopenia status in patients with CKD, is shown in Figure 2. The Kaplan-Meier analysis revealed that the risk of hospitalization was gradually increased in presarcopenic and sarcopenic patients, compared to that in the normal group (log-rank test: p < 0.001). In the univariate Cox proportional analysis, age, the administration or lack of dialysis, the presence of cardiac disease, serum albumin levels, serum creatinine levels and sarcopenia were significant predictors of all-cause hospitalizations in CKD patients (Table 4). After adjustments for all of the possibly related covariables (age, sex, dialysis or no dialysis, the presence of comorbidities, BMI and serum levels of albumin, creatinine and 25(OH)D; model 3), the multivariate Cox proportional hazards models demonstrated that the risk of hospitalization was significantly higher for patients with presarcopenia (hazard ratio, 2.48; 95% confidence interval, 1.180–5.230) and much higher for patients with sarcopenia compared to that in the normal group (hazard ratio, 9.11; 95% confidence interval, 2.295–25.182, Table 5).

**Figure 2.**
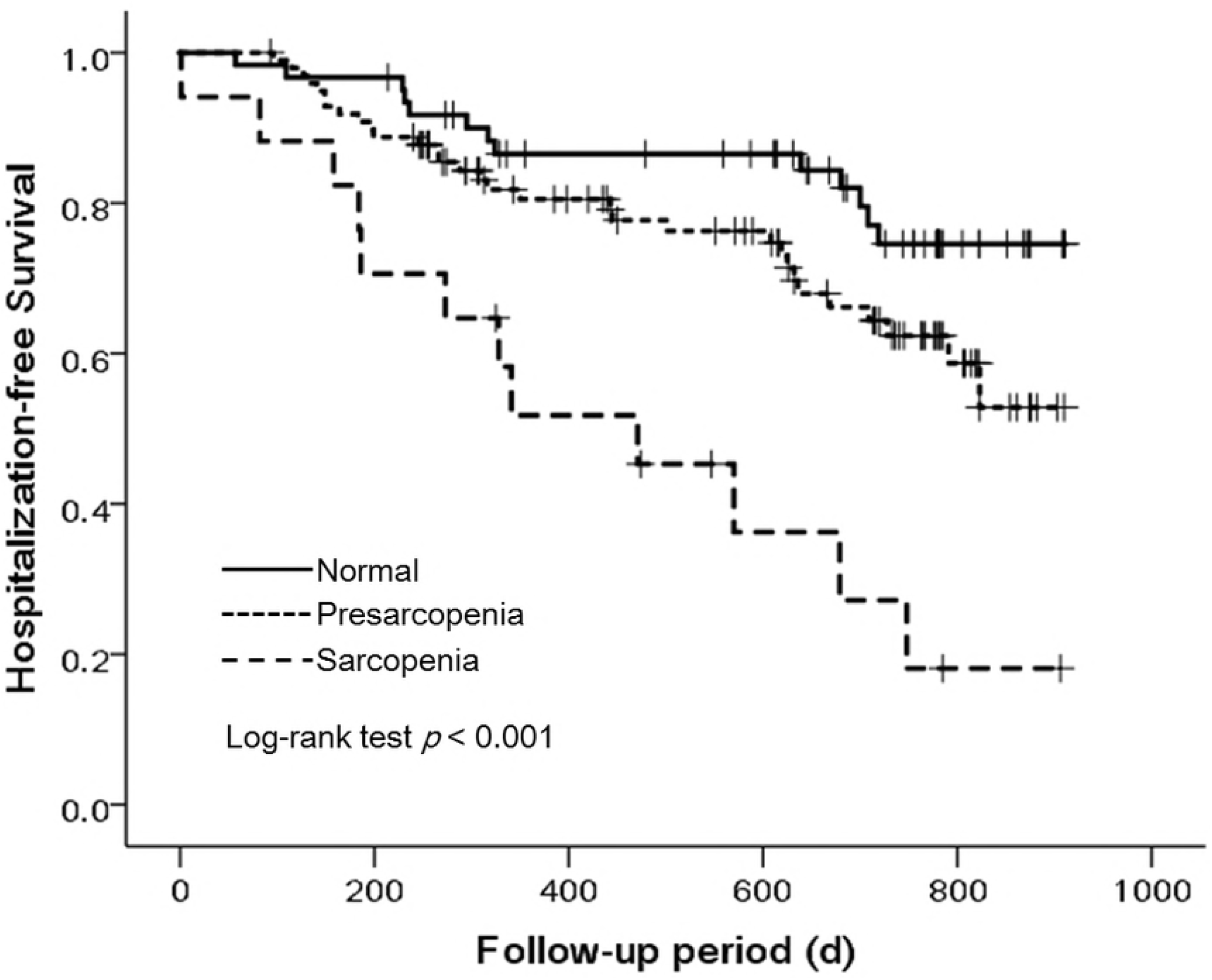
Kaplan-Meier estimates of hospitalization-free survival probabilities of the patients with chronic kidney disease in relation to sarcopenia status and categorized by the ASM/wt index. *P*, P-value.

**Table 4.** Univariate Cox proportional models for hospitalization. in patients with chronic kidney disease.

| Characteristics | HR (95%CI) | P-value |
| --- | --- | --- |
| Age (year) | 1.03 (1.005 – 1.047) | 0.016 |
| Sex (female/male) | 0.69 (0.408 – 1.171) | 0.170 |
| Dialysis (no/yes) | 1.86 (1.084 – 3.188) | 0.024 |
| Diabetes (absent/present) | 1.41 (0.833 – 2.384) | 0.200 |
| Hypertension (absent/present) | 0.87 (0.447 – 1.677) | 0.668 |
| Cerebrovascular (absent/present) | 0.98 (0.442 – 2.158) | 0.954 |
| Cardiac (absent/present) | 3.22 (1.800 – 5.756) | <0.001 |
| BMI (kg/m<sup>2</sup>) | 0.95 (0.895 – 1.006) | 0.076 |
| Albumin (g/dL) | 0.40 (0.237 – 0.675) | 0.001 |
| Creatinine (mg/dL) | 1.08 (1.019 – 1.144) | 0.009 |
| 25(OH)D (ng/mL) | 0.98 (0.957 – 1.008) | 0.181 |
| Presarcopenia (vs normal) | 1.83 (0.955 – 3.498) | 0.069 |
| Sarcopenia (vs normal) | 5.38 (2.438 – 11.857) | <0.001 |

25(OH)D, 25-hydroxyvitamin D; BMI, body mass index; CI, confidence interval.

**Table 5.** Multivariate Cox proportional models for hospitalization in patients with chronic kidney disease.

| Characteristics | HR (95%CI) | P-value |
| --- | --- | --- |
| **Model 1** |  |  |
| Age (year) | 1.02 (0.995 – 1.039) | 0.142 |
| Sex (female/male) | 0.77 (0.449 – 1.327) | 0.349 |
| Presarcopenia (vs normal) | 1.67 (0.864 – 3.240) | 0.127 |
| Sarcopenia (vs normal) | 4.15 (1.778 – 9.679) | 0.001 |
| **Model 2** |  |  |
| Age (year) | 1.01 (0.986 – 1.032) | 0.449 |
| Sex (female/male) | 0.80 (0.450 – 1.408) | 0.433 |
| Dialysis (no/yes) | 1.32 (0.740 – 2.345) | 0.349 |
| Diabetes (absent/present) | 1.03 (1.027 – 0.569) | 0.930 |
| Albumin (g/dL) | 0.39 (0.190 – 0.794) | 0.010 |
| Presarcopenia (vs normal) | 1.82 (0.917 – 3.627) | 0.087 |
| Sarcopenia (vs normal) | 4.18 (1.702 – 10.253) | 0.002 |
| **Model 3** |  |  |
| Age (year) | 1.01 (0.982 – 1.039) | 0.500 |
| Sex (female/male) | 0.76 (0.396 – 1.449) | 0.401 |
| Dialysis (no/yes) | 0.20 (0.058 – 0.702) | 0.012 |
| Diabetes (absent/present) | 1.27 (0.629 – 2.510) | 0.518 |
| Hypertension (absent/present) | 1.22 (0.506 – 2.950) | 0.656 |
| Cerebrovascular (absent/present) | 1.26 (0.465 – 3.419) | 0.649 |
| Cardiac (absent/present) | 3.15 (1.527 – 6.514) | 0.002 |
| BMI (kg/m<sup>2</sup>) | 0.97 (0.903 – 1.037) | 1.037 |
| Albumin (g/dL) | 0.39 (0.166 – 0.906) | 0.029 |
| Creatinine (mg/dL) | 1.23 (1.072 – 1.417) | 0.003 |
| 25(OH)D (ng/mL) | 0.99 (0.959 – 1.019) | 0.452 |
| Presarcopenia (vs normal) | 2.48 (1.180 – 5.230) | 0.017 |
| Sarcopenia (vs normal) | 9.11 (2.295 – 25.182) | <0.001 |

25(OH)D, 25-hydroxyvitamin D; BMI, body mass index; CI, confidence interval.

## Discussion

In our study, skeletal muscle mass was measured via a bioimpedance analysis, and the handgrip test was used in all of the participants. In terms of the adjusted ASM, the ASM/wt index showed the most favorable correlations with chronological age, muscle strength, insulin resistance, and frailty scores compared to those of other indices in these populations. When patients were categorized into normal, presarcopenia, or sarcopenia groups, the patients with presarcopenia and sarcopenia showed poor hospitalization-free survivals.

In 1974, Floyd et al. documented uremic myopathy in CKD patients (26), and Clyne suggested that the prevalence of uremic myopathy was as high as 50% in dialysis patients (27). In cross-sectional studies comparing muscle biopsy samples between age-matched controls and ESRD patients, fiber atrophy and reduced capillary density were observed (28), and a significant decrease in the mean diameters of both type I and II fiber types (29) was reported. Additionally, decreased muscle function has been suggested in chronic dialysis and predialysis patients with CKD by recording electrically stimulated muscle contractions (27, 30). Foley et al. reported that there was a significant association between the increased prevalence of sarcopenia and the decline in glomerular filtration rate (31). Similarly, in a more recent study in Korea with 11,625 subjects aged 40 years or older, the prevalence of low adjusted ASM (ASM/wt) was associated with CKD stage in men (32). In a study that utilized a recent definition of sarcopenia, which was represented by the simultaneous incidence of low muscle mass and decreased power, the prevalence of sarcopenia was 5.9-9.8%, depending on the muscle mass assessment method used in the predialysis CKD patients (33). Lamarca et al. investigated sarcopenia in elderly maintenance hemodialysis patients and found that the prevalence of muscle weakness that was measured by handgrips was 85%, but the prevalence of decreased muscle mass varied from 4-73.5%, depending on the method that was used and the cutoff value that was applied (34).

One of the challenges in studies of sarcopenia is to determine how best to measure the amount of muscle. However, the European Working Group on Sarcopenia in Older People, as well as the AWGS, recently established a clear criterion for the diagnosis of sarcopenia (35, 36). The guideline recommends the use of a bioimpedence analysis, dual-energy X-ray absorptiometry, computed tomography, and magnetic resonance imaging to measure the skeletal muscle mass. Bioimpedance analysis, which was used in our study, is a popular method for estimating body composition and has the merits of being low-cost, easy-to-use and portable. Furthermore, the bioimpedance analysis results, under standard conditions, have been found to correlate well with MRI predictions, and the prediction equations have been validated for various populations (37). In our data, the combined prevalence of sarcopenia and presarcopenia, through the use of adjusted ASM that was measured by bioimpedance analysis, was approximately 60% (Table 2). Finally, approximately 40% of the subjects had normal muscle mass and strength. Our data showed that muscle failure is common in CKD patients, which is consistent with other epidemiologic studies of sarcopenia or low muscle mass in that population (11-13). In comparison with the results from healthy Korean women who were older than 65 years old, 21% of them were classified into either presarcopenia or sarcopenia groups (38).

Sarcopenia does not involve only muscle mass or strength but also involves basal metabolic rates (39). Muscles are the largest repository of protein, the most important site for the storage of glucose and the major critical consumer of energy. Thus, sarcopenia contributes to decreased energy demands of the human body (39). Of course, sarcopenia, including reduced muscle mass, has negative consequences for human health. Muscle weakness, which is one component of sarcopenia, has been reported to be associated with poor health outcomes, such as mortality, hospitalization, and disabilities in predialysis and dialysis patients (40). Pereira et al. suggested that predialysis patients with sarcopenia had higher mortality rates (33). In our study, both sarcopenia and presarcopenia were predictive of hospitalization-free survival among CKD outpatients, including patients in the predialysis and dialysis stages (Figure 2).

Exercise interventions could be the primary treatment options for sarcopenia. Indeed, most exercise trials have shown improved muscle strength, physical performance and muscle mass in community-dwelling, elderly individuals, although these individuals were not recruited based on their sarcopenic status (41-44). Additionally, essential amino acid and β-hydroxy β-methylbutyric acid (HMB, which is a bioactive metabolite of leucine) supplements seem to have some effects on muscle mass and muscle function; however, these effects need to be confirmed in larger trials (45-48). Vitamin D, testosterone, or myostatin inhibitors have also been proposed as potential drugs for the treatment of muscle failure (49-51). Remarkably, there are several studies that support the idea that exercise may enhance muscle strength in CKD patients (52). Howden et al. reported that 12-month exercise and lifestyle interventions improved 6-minute walk distances in patients with CKD stages ranging from 3 to 4 (53). Greenwood et al. showed that the completion of a 12-week exercise rehabilitation regimen enhanced physical performance and reduced cardiovascular morbidity (54). However, the effects of exercise training on sarcopenia are unclear in patients with CKD. To confirm these effects, it is necessary to identify the prevalence of sarcopenia, according to the recently suggested definition and to examine its relationship with clinical outcomes among the CKD patient population.

There were several limitations of our study. First, this was a small, single-center study that had a relatively short observation period. However, this study had several strengths. We used a definition of sarcopenia that included the two components of low muscle mass and muscle strength. Furthermore, we used a direct physical function test to estimate muscle strength. We compared the prevalence of sarcopenia by using three different muscle indices and explored the most appropriate muscle indices in CKD patients, which may reflect their chronological age, muscle power, metabolic derangement, and frailty. Lastly, we examined whether the selected muscle index was a good predictor for clinical prognoses in our subjects.

In conclusion, our findings showed that sarcopenia and presarcopenia can be useful for predicting hospitalization in CKD outpatients. Future studies on sarcopenia may provide new methods for gaining insights into the disease and for improving their prognoses. Therefore, we should recognize the sarcopenic and presarcopenic statuses of patients as risk factors for poor clinical outcomes and proceed with further research on the relationship between these risk factors and disease status.

## Ethic statement

This study was approved by an independent Ethics Committee at the CHA Bundang Medical Center, and written informed consent was obtained from each of the patients.

## Author Contribution Statement

H. Y. Jeong and W. Ahn wrote the first manuscript text; J. C. Kim and Y. B. Choi prepared Figures 1-2; J Kim, H. H. Jun and S. Lee prepared Table 1-5; D. H. Yang and J. Oh analyzed and interpreted the data; J. Bae and S-Y Lee designed the study and approved the final vision of the manuscript and figures to be submitted and published.

